# Comparable phosphene size and contrast properties, but differing detection reliability for optogenetic versus electrical stimulation of macaque V1

**DOI:** 10.64898/2026.09.03.749087

**Authors:** Kumari Liza, Marta Falkowska-Kisiel, Jaime Cadena-Valencia, Jennifer Greilsamer, Florian Lanz, Marcus Haag, Alessandra Bergadano, Diego Ghezzi, Michael C. Schmid

## Abstract

Cortical visual prostheses aim to restore vision by stimulating primary visual cortex (V1) to evoke artificial percepts (phosphenes). Electrical stimulation has long served this purpose, and optogenetic approaches promise higher spatial resolution via cell-type specificity. Recent investigations have demonstrated localized behavioral responses to optogenetic V1 stimulation (1–3), but the applied perimetry paradigms cannot dissociate a genuine percept from a reflexive oculomotor response, leaving the perceptual character of optogenetic phosphenes unresolved.

To address this gap, two macaques received optogenetic V1 injections guided by laminar electrophysiology and performed detection and discrimination tasks under interleaved electrical, optogenetic, and visual stimulation, enabling direct within-subject comparison.

Electrical stimulation reliably evoked detectable percepts (positive d′ in 100% of sessions), with thresholds scaling with current, resembling visual contrast-response functions. Optogenetic stimulation was far less reliable, with positive d′ in only ∼50% of sessions, despite confirmed V1 activation. Despite this reliability difference, optogenetically evoked percepts closely resembled electrically and visually evoked percepts in perceptual character: perceived size (∼1° of visual angle) was comparable across conditions and scaled similarly with eccentricity. Similarly, perceived contrast equivalents (5%-11% visual-contrast equivalent) were broadly comparable across stimulation approaches.

These findings indicate that successful optogenetic activation engages the same perceptual encoding mechanisms as electrical and natural vision, but that reliable detection may additionally depend on circuit-level mechanisms not consistently engaged by current optogenetic parameters. The comparable percept size further suggests optogenetic stimulation may not confer the enhanced spatial resolution often proposed as its key advantage.

**Significance statement:** Brain implants that stimulate visual cortex could restore vision to people blinded by damage to the eye or optic nerve, evoking flashes of light called phosphenes. Electrical stimulation, the currently used approach, activates tissue broadly; optogenetic stimulation, which uses light to activate genetically targeted neurons, has been proposed as a more precise alternative, but whether it produces a genuine visual sensation was unclear. Using monkeys trained to report artificially evoked flashes, we show that optogenetic stimulation evokes sensations resembling electrical stimulation in size and brightness but is detected far less consistently. These results show optogenetic stimulation engages the same perceptual pathways as natural vision, but does not yet deliver its assumed resolution advantage, informing the design of future optical visual prostheses.

## Introduction

Cortical visual prostheses aim to restore vision by directly stimulating neurons in the visual cortex, thereby bypassing damaged retinal and optic nerve pathways. Although electrical stimulation of the human visual cortex has been known to evoke artificial visual percepts, commonly referred to as phosphenes for nearly a century, generating phosphenes with sufficient fidelity for functional, long-term vision remains a major challenge. Early studies in blind and sighted human participants using surface electrodes demonstrated that electrical stimulation of V1 can reliably evoke localized phosphenes whose position reflects the underlying retinotopic map (4–6). More recently, intracortical microelectrode arrays have enabled stimulation with substantially improved spatial precision and lower currents, effectively dropping thresholds from the milliampere range required for surface stimulation down to the microampere range (often below 25 uA) (7, 8). Human and non-human primate studies have shown that phosphene size generally increases with eccentricity and with stimulation intensity, although it rapidly reaches a saturation point at moderate current amplitudes (7). Meanwhile, brightness can be actively modulated by stimulation current and pulse parameters, such as pulse frequency and duration. However, despite considerable technological progress, current tissue penetrating devices carry risks regarding long-term tissue health (10–12). Furthermore, electrical microstimulation lacks cellular specificity because it preferentially activates highly excitable myelinated axons, including axons of passage belonging to neurons whose cell bodies lie well outside the nominal stimulation radius. It activates neurons across a substantially larger and more heterogeneous volume of tissue than rheobase/chronaxie-based current-distance estimates alone would suggest (13). Behaviorally discriminable phosphene locations in the visual field require electrode separations in macaque V1 on the order of 1.6 mm (14, 15), and fMRI measurements of microstimulation-evoked activity reveal an even broader, trans-synaptically mediated spread that extends into retinotopically connected but spatially remote visual areas (14, 15). This indiscriminate, spatially extensive recruitment produces neural activity patterns that differ substantially from those generated during natural vision.

Optogenetic targeting of V1 has been proposed as an alternative to electrical stimulation. It offers genetically defined cell-type specificity for targeting perceptually relevant circuit elements (16). Computational models further suggest it can achieve spatial resolution on the order of 100 μm (17) — one to two orders of magnitude finer than the behaviorally and hemodynamically defined spread of electrical microstimulation described above. In rodents, where optogenetics has revolutionized circuit analysis, the method has been successfully applied to visual cortex, so that mice can detect and discriminate between varying optogenetic stimulation patterns (18). In macaques, a direct within-study comparison at the level of the LGN-V1 circuit found that optogenetic activation of koniocellular LGN neurons and electrical microstimulation of the same LGN layers produced the same supra-granular V1 activation pattern, indicating comparable capacities of both methods to isolate this thalamocortical pathway (19). Consistent with this convergence, we separately showed that optogenetic stimulation of V1 itself drives local V1 activity as well as downstream extrastriate BOLD fMRI activity in areas V2/V3, V4, MT, and FEF, in a pattern qualitatively similar to that elicited by electrical microstimulation of V1 (19, 20). Behaviorally, two studies demonstrated eye movements towards retinotopic locations targeted by optogenetic stimulation of V1 (1, 2). Conversely, optogenetic stimulation targeting V1 interneurons was shown to disrupt vision guided behavior (21). However, attempts to delineate the perceptual quality of optogenetic V1 stimulation have been challenging and remain far less advanced than our knowledge about phosphene quality elicited by electrical V1 stimulation. Here, we directly compare electrical and optogenetic stimulation of macaque V1 using a combination of psychophysics and laminar electrophysiology. We quantify the size and brightness of optogenetically induced visual percepts and directly compare them to those obtained from electrical stimulation.

## Materials and Methods

### Subjects

Two adult male rhesus macaques (Macaca mulatta) were used for this study (Monkey G: 8 kg and Monkey C: 10 kg). All animal procedures and experimentations were reviewed and approved by the cantonal animal ethics board and veterinary office and performed in accordance with the Swiss animal protection law. The monkeys were housed at the University of Fribourg in groups of 2-3 animals in 45 m^3^ rooms, with opportunities to climb, forage, engage with enrichment tools and with daily access to an outside area with natural sunlight. The monkeys had unlimited access to water. Food in the form of pellets and vegetables was typically delivered after an experimental session in accordance with animal’s nutritional requirements and welfare needs. Motivation for visual task engagement was reinforced by providing preferred sweet food such as fruits and fluids such as juice during experimental testing sessions in the laboratory. Monkeys were first trained to perform visual tasks under minimal restraint using the chinrest approach (22). Titanium headpost and chamber implants (Rogue Research Inc., Montreal, Canada) were individually fitted based on CT and MRI scans (23). Access to V1 was provided using a 13 mm wide craniotomy over the right occipital cortex. All surgical procedures were performed under refined anesthesia protocols (24).

### Visual stimulation and behavioral paradigms

Visual stimuli were generated in MWorks v0.10 (https://mworks.github.io) with custom-written scripts, running on an iMac under macOS Mojave. Stimuli were rear-projected by a PROPixx projector (VPixx Technologies Inc., Saint-Bruno, QC, Canada) onto a translucent projection screen (1308 × 737 mm, VPixx Technologies Inc.) at 1920 × 1080 pixels and a refresh rate of 120 Hz. All stimuli were rendered with a uniform grey background of 45 cd/m². Animals were head-fixed with their eyes at 57 cm distant from the screen. Gaze position of the left eye was recorded at 1000 Hz with an infrared eye tracker (EyeLink 1000, SR Research Ltd., Ottawa, ON, Canada).

### Receptive field mapping

To map receptive fields (RFs), animals fixated on a central fixation point (0.2 dva central dot within a 1–1.5 dva window) while wedges or annuli were presented on screen, as previously described (25). Wedges with an opening angle of 5° were presented sequentially in orientations spanning from 180° to 270° in 2.5° steps (50% overlap). Annuli eccentricities ranged from 1° to 9°, with thickness matched to the cross-sectional width of a wedge at the corresponding eccentricity and spacing chosen to yield 50% overlap between successive annuli. Both annuli and wedges consisted of a drifting grating with a spatial frequency of 4 and a speed of 3 deg/s. Each trial presented 3 to 10 stimuli were displayed (random geometric distribution), each shown for 100 ms. Only trials where fixation maintained throughout were used for analysis and animals were rewarded for holding fixation across the full sequence of a trial.

### Detection task

Detection was tested in blocks. Each trial began with a central fixation dot, fixation had to be held within a 1–1.5 dva window for 500 ms (Monkey G) or 700 ms (Monkey C) before stimulus or stimulation onset with variability of 0-300 ms (random geometric distribution). On visual trials (50%), a 1° square of varying contrast (0.5-40%, Weber contrast) was presented at the V1 RF location (lower left quadrant) along with two response targets in the upper quadrants. On the remaining catch trials (50%), only two response targets were presented. Animals were required to make a saccade to the upper left target on visual target or the upper right target on catch trials. In phosphene detection block (Fig. 1D), instead of a visual stimulus, V1 was either electrically (train duration: 50 ms) or optogenetically (train duration: 200 ms) stimulated in the RF location. Similar to the visual detection block, half of the trials included stimulation and the remaining 50% were catch trials where no stimulation was sent. Animals were required to respond to detection of a phosphene by a saccade to the upper left target or to the upper right target on catch or undetected trials. Trials with saccades made within 50 ms of stimulus or stimulation onset were excluded as early reflexive responses. Animals had upto 500 ms to respond. Trials were aborted if fixation broke before the response. Each correct trial was rewarded with juice.

**Fig. 1.**
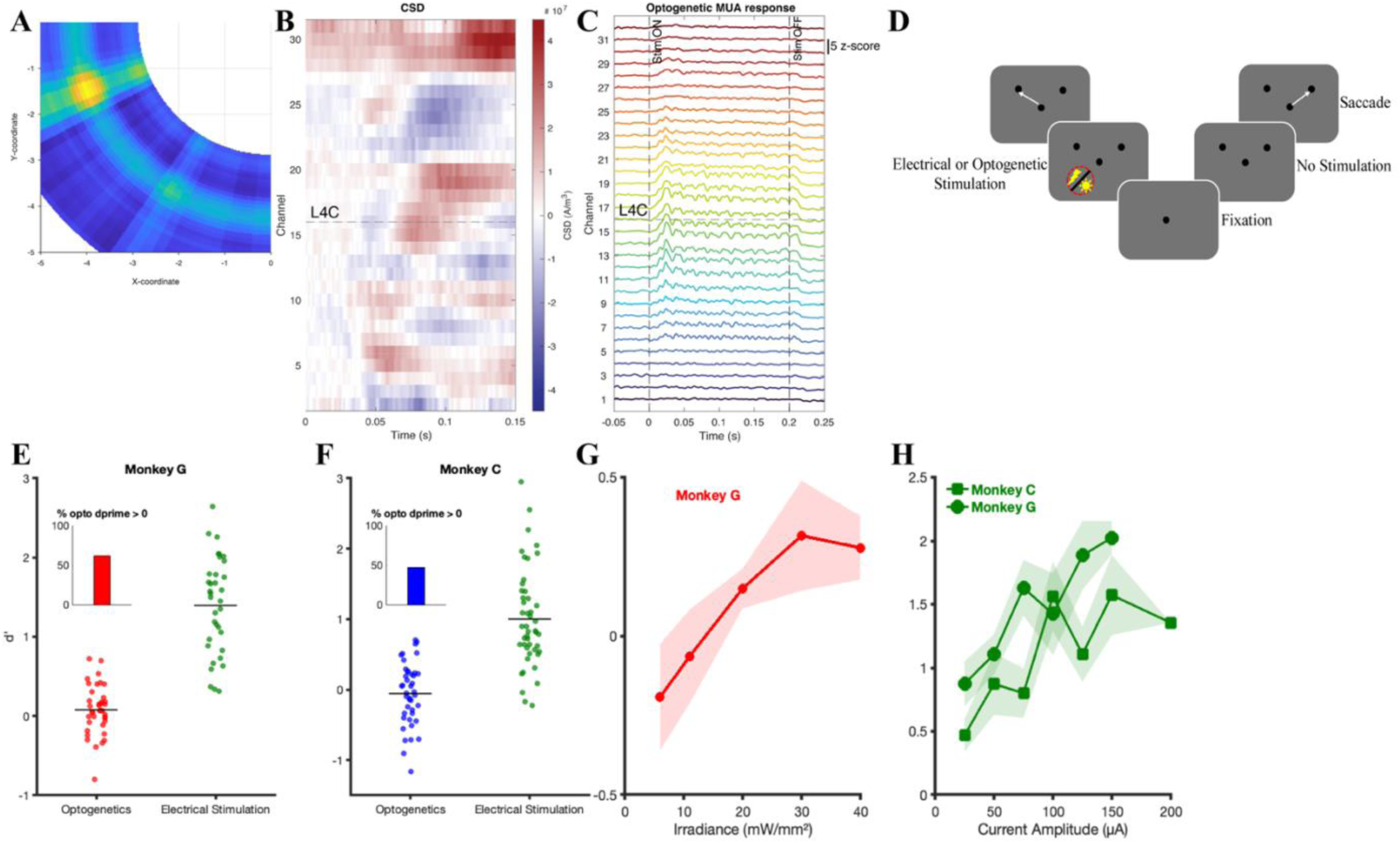
Phosphene detection task with electrical and optogenetic stimulation. (A) RF location for an example recording site in Monkey G during an electrical microstimulation session. (B) CSD profile to visual stimulation for an example recording site in Monkey G during an optogenetic session. (C) MUA activity from the same session, evoked by epidural red LED stimulation. (D) Schematic of the phosphene detection task under electrical or optogenetic stimulation. (E and F) d’ of all sessions of optogenetic and electrical stimulation for Monkey G (E) and Monkey C (F). Inset bar graph represents the proportion of sessions with d’>0; ∼67% for Monkey G and ∼50% for Monkey C. (G) d’ as a function of irradiance for optogenetic (red LED) stimulation in Monkey G. (H) d’ as a function of current amplitude for electrical microstimulation in Monkey G and Monkey C.

### Discrimination task

In discrimination tasks, 50% of trials were visual trials and the remaining 50% were stimulation trials. The minimum fixation duration was 500 ms with stimuli or stimulation onset variability of 0-300 ms (random geometric distribution). On visual trials, two discs were presented simultaneously -- one at the RF (lower left quadrant) and one diagonally opposite (upper right quadrant). On stimulation trials, the RF location was electrically or optogenetically stimulated, while a disc was presented simultaneously in the upper right quadrant. The stimulation trigger was delayed ∼30 ms to approximate retina to V1 conduction delay of a visual stimulus. Animals were trained to saccade towards the stimulus or phosphene that was either brighter or larger.

On visual trials (Fig. 2A & 4A), the RF disc served as the ‘standard’ (fixed contrast or size) and the disc presented diagonally opposite served as the ‘probe’ (variable contrast or size). Similarly, on stimulation trials (Fig. 2D & 4D), stimulation strength (current amplitude or irradiance) was held constant within a block while probe contrast or size varied. As above, responses within 50 ms of stimulus or stimulation onset were excluded and animals had 500 ms to respond. On stimulation trials, rewards were either distributed randomly (50% of the time), irrespective of choice or based on fixed rule in a block (probe value relative to median of its range was taken as a threshold for rewarding) with results remaining consistent across both rewarding methods. Independent experiments (Fig. S2) confirmed that the reward mode had no influence on the obtained results.

**Fig. 2.**
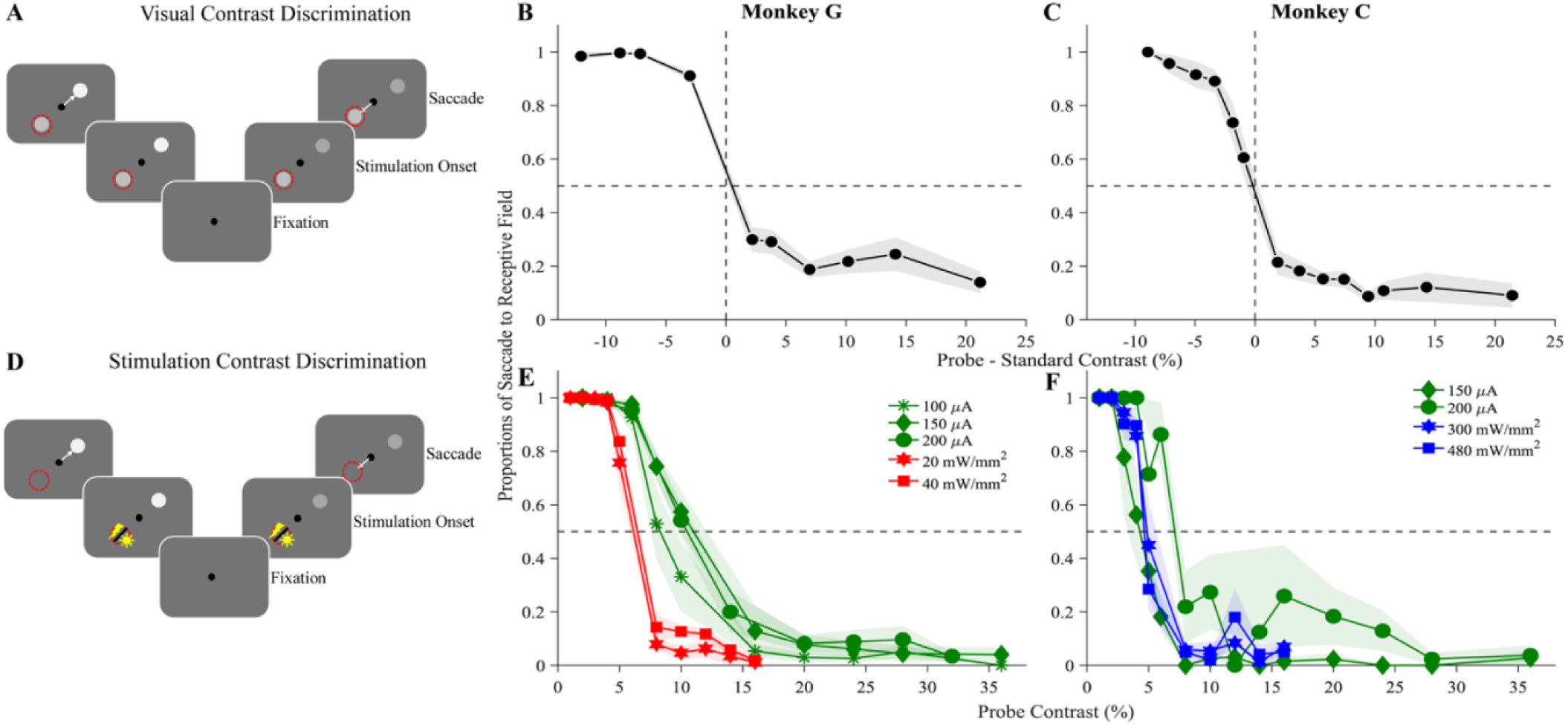
Contrast discrimination with visual, electrical and optogenetic stimulation. (A) Schematic of the visual contrast discrimination task. On each trial, the animal saccades to the brighter stimulus either in the RF or outside the RF. (B and C) Proportion of saccades towards the RF location as a function of percentage contrast difference between probe and standard stimuli for Monkey G (B) and Monkey C (C). (D) Schematic of the stimulation contrast discrimination task. The standard visual stimulus was replaced by electrical or optogenetic stimulation delivered at the RF location. (E and F) Proportion of saccades towards the RF location as a function of probe contrast for Monkey G (E) and Monkey C (F). Green traces show electrical stimulation trials at current amplitudes of 100, 150 and 200 µA (E) and 150 and 200 µA (F); red/blue traces show optogenetic stimulation trials at irradiances of 20 and 40 mW/mm² (E, Monkey G) and 300 and 480 mW/mm² (F, Monkey C).

### Viral Injections

Injections into V1 were performed in the awake behaving animal via the recording chamber implant. For optogenetic activation, Monkey C was injected with AAV9-hSyn-hChR2(H134R)-EYFP (#26973; Addgene) viral vector. Monkey G received an injection of AAV9-CamKIIa-ChrimsonR-mScarlet-KV2.1 (#124651, Addgene). Injections were administered using a 24-channel laminar electrode array (Plexon Inc.) featuring 3 fluid capillaries at distances of 1 mm, 1.7 mm, and 2.4 mm from the electrode tip. Monkey C was in total injected at 4 sites with an average of 6.5 µL per site at the rate of 43.3 nL/min respectively. Monkey G received ChrimsonR injections at 3 sites (mean volume 22.6 µL/site) delivered at 240 nL/min.

### Optogenetic stimulation

Optogenetic experiments commenced six weeks after viral injections. In Monkey C, blue light stimulation from a 473 nm Omicron LuxX diode laser was delivered into the neuropil through a Plexon optrode at 40 Hz for 200 ms, with irradiances ranging from 100 to 500 mW/mm². Irradiance was measured prior to each recording session at the tip of the optrode using a sphere sensor and a digital power meter (Thorlabs). Neuropil stimulation was performed using a 24- or 32-channel Plexon S-probe incorporating a 50 µm optic fiber (0.66 NA) positioned between channels 15 and 16. The probe had an interelectrode spacing of either 75 or 100 µm, depending on the number of recording channels, and the probe shaft served as the reference electrode for electrophysiological recordings. In Monkey G, red light stimulation was delivered epidurally using a 653 nm LED (Prizmatix) coupled to an optic fiber (1500 µm diameter, 0.5 NA, Thorlabs) at 80 Hz for 200 ms. The irradiance ranged from 5 to 40 mW/mm².

### Electrical microstimulation

Electrical stimulation was delivered using a CereStim^TM^ R96 stimulator (Blackrock Microsystems). Anodic-first biphasic charge balanced pulses of 25-200 μA current amplitude with pulse width of 200 μs, 60 μs interphase interval, 200 Hz frequency and stimulation duration of 50-200 ms were delivered via monopolar tungsten electrodes (diameter: 280 μm with insulation, FHC Inc.). All recordings were performed at depths of 0.2 to 1.5 mm from the dura mater. Electrode impedance ranged from 400kΩ to 1 MΩ. A silver wire was placed over the dura mater served as the reference.

### Electrophysiological Recordings and Analyses

To advance through the dura mater, the S-probe and FHC electrode were placed in a stainless-steel guide tube and advanced into V1 using a hydraulic micromanipulator (MO-97A, Narishige). Neurophysiological signals were acquired at a 30 kHz sampling rate using a Blackrock recording system (Blackrock Microsystems, Inc.; 32-CH Omnetics Headstage connected to a front-end amplifier; a 32-CH FHC to Omnetics adaptor was used for the tungsten electrode). Data analyses were done in MATLAB with custom-written scripts using the Fieldtrip toolbox (26). Multi-unit activity (MUA) signals were extracted by applying a high-pass filter between 300 Hz and 12 kHz, followed by full-wave rectification (27). Local field potential (LFP) signals were extracted by low-pass filtering the raw signal at 150 Hz. For all subsequent analyses, both MUA and LFP signals were downsampled to 3000 Hz.

At the start of each recording session, we mapped the receptive field (RF) of the recording site by measuring the average multi-unit activity (MUA) response 50–100 ms after stimulus onset across all stimulus locations (Fig. 1A). For sessions with S-probe recording spanning the full depth of the V1 laminar profile, we additionally computed the current source density (CSD) profile from responses to a 1-degree square presented in the RF location during fixation, using the sink with the shortest response latency (typically layer 4C) to confirm probe placement (Monkey G, Fig. 1B & Monkey C, Fig. S1A). At each site, we further verified successful stimulation by confirming a clear MUA response to the optogenetic (LED or laser) stimulation (Monkey G, Fig. 1C & Monkey C, Fig. S1B).

Sensitivity or d’ was computed using signal detection theory (28). To avoid infinite values arising from hit or false alarm rates of 0 or 1, we applied the log linear correction (29), adding 0.5 to the hit and false alarm counts and 1 to the number of visual and catch trials before computing proportions.

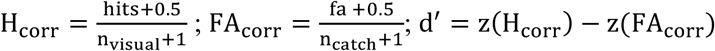

Here, H_corr_ , hits and n_visual_ are corrected hit rate, number of hits in visual trials and number of visual trials respectively. FA_corr_, fa and n_catch_ are corrected false alarm rate, number of false alarms in catch trials and number of catch trials respectively. z is the inverse of the standard normal cumulative distribution function.

## Results

### Low phosphene detectability for optogenetic compared to electrical stimulation

We first confirmed that electrophysiological recording and stimulation sites were in V1 using RF mapping (Fig. 1A, example from an electrical stimulation session in Monkey G) and, for laminar recordings, using CSD analysis in response to visual stimulation (see Methods). Fig. 1B–C shows an example optogenetic session in Monkey G, with the CSD profile to visual stimulation confirming the laminar recording spanned V1 (Fig. 1B; Monkey C, Fig. S1A) and a clear LED-evoked MUA response, concentrated in the input (granular) layer, confirming successful optogenetic activation at the site (Fig. 1C; Monkey C, Fig. S1B: Laser-evoked MUA response).

We next tested whether electrical and optogenetic stimulation were equally effective at evoking detectable phosphenes. Detection differed markedly between stimulation modalities. Electrical stimulation reliably evoked detectable phosphenes: computing d’ for the phosphene detection task (Fig. 1D) across sessions in Monkey G (Fig. 1E) and Monkey C (Fig. 1F), positive d’ was observed across all current amplitudes tested and performance increased consistently with current amplitude (Fig. 1H). Optogenetic stimulation was considerably less consistent: positive d’ across all irradiances was observed in 63% and 48% of sessions in Monkey G and C, respectively. In a subset of Monkey G session with at least three irradiance levels tested under matched stimulation parameters, d’ increased with irradiance (Fig. 1G). No comparable trend was observed in Monkey C. Across all irradiances tested, highest d’ achieved with optogenetic stimulation was comparable to d’ observed at lowest current amplitude used for electrical stimulation.

### Electrical and optogenetic stimulation induce phosphenes of comparable contrast

In visual contrast discrimination trials (Fig. 2A), we measured the proportion of saccades directed to RF (where the standard stimulus was presented) as a function of contrast difference between probe and standard stimuli, in Monkey G (Fig. 2B) and Monkey C (Fig. 2C). Results from statistical assessment are summarized in Table 1. As expected, when the probe contrast was lower than that of the standard, animals consistently saccaded towards the RF; as probe contrast increased beyond that of the standard, the proportion of saccades to the RF decreased. To quantify the perceived contrast of induced phosphenes, we replaced the standard stimulus with electrical or optogenetic stimulation delivered at the RF in Monkey G (Fig. 2E) and Monkey C (Fig. 2F). As in the visual task, at low probe contrast, animals consistently saccaded towards the RF (i.e. towards stimulation); at higher probe contrast, they switched to saccades towards the probe stimulus (i.e. away stimulation).

**Table 1:** Comparison of electrical and optogenetic stimulation across behavioral measures. For each measure, sessions were pooled across all current amplitudes (electrical) or irradiances (optogenetic), and the two pooled distribution were compared using a two-sided Wilcoxson rank sum test. n_Elec and n_Opto denote the number of sessions contributing to electrical microstimulation and optogenetic pooled conditions respectively. Each measure was tested independently, and p-values are not corrected for multiple comparisons across measures.

| Task | Measure | Monkey G |  |  |  | Monkey C |  |  |  |
| --- | --- | --- | --- | --- | --- | --- | --- | --- | --- |
|  |  | *n_Elec* | *n_Opto* | *pval* | *Effect size (Elec-Opto)* | *n_Elec* | *n_Opto* | *pval* | *Effect size (Elec-Opto)* |
| Contrast Discrimination | Threshold | 15 | 13 | <0.001 | 0.98 | 10 | 8 | 0.46 | 0.23 |
|  | RT (towards stim.) | 15 | 13 | 0.01 | -0.58 | 10 | 8 | <0.001 | -0.95 |
|  | RT (away stim.) | 15 | 13 | 0.013 | -0.56 | 10 | 8 | 0.63 | -0.15 |
|  | End Point Scatter (towards stim.) | 15 | 13 | 0.019 | -0.53 | 10 | 8 | <0.001 | -0.98 |
|  | End Point Scatter (away stim.) | 15 | 13 | <0.001 | -0.99 | 10 | 8 | 0.003 | -0.80 |
| Size Discrimination | Threshold | 15 | 16 | 0.005 | -0.60 | 10 | 10 | 0.173 | -0.37 |
|  | RT (towards stim.) | 15 | 16 | <0.001 | -0.78 | 10 | 10 | 0.385 | -0.24 |
|  | RT (away stim.) | 15 | 16 | <0.001 | -0.94 | 10 | 10 | 0.121 | -0.42 |
|  | End Point Scatter (towards stim.) | 15 | 16 | <0.001 | -0.78 | 10 | 10 | 0.623 | 0.14 |
|  | End Point Scatter (away stim.) | 15 | 16 | <0.001 | -0.94 | 10 | 10 | 0.791 | 0.08 |

The phosphene contrast threshold was defined as the probe contrast at which the animal was equally likely (0.5 proportion) to saccade towards stimulation or towards probe stimulus. This threshold represents the perceived contrast of the induced phosphene, expressed in equivalent visual contrast. In Monkey G, optogenetic stimulation elicited phosphenes with an effective contrast of approximately 6%, which did not change with increasing irradiance. Conversely, electrical stimulation in Monkey G elicited phosphenes with significantly higher perceived contrast (9% to 11%), which increased incrementally as the current amplitude was raised (Fig. 3A). This difference in perceived phosphene in Monkey G could be due to epidural optogenetic stimulation compared to intracortical electrical stimulation. In Monkey C, the effective contrast of phosphenes was found to be comparable (4.5 to 7.5%) between electrical and optogenetic stimulation (Fig. 3B). We also examined behavioral responses associated with phosphene detection. In both monkeys, increasing the current amplitude of electrical stimulation and irradiance produced a modest reduction in saccadic reaction time for saccades towards stimulation condition, suggesting a slight increase in phosphene saliency in both Monkey G (Fig. 3C) and Monkey C (Fig. 3E). Electrical stimulation trials had significantly lower reaction times compared to optogenetic trials (Fig. 3C & Fig 3E). Compared to the towards stimulation condition, the reaction times for away stimulation condition (i.e. towards probe stimulus) were substantially shorter across both stimulation techniques in Monkey G (Fig. 3D) and Monkey C (Fig. 3F). To assess the spatial precision of the induced phosphene, we calculated endpoint scatter (the standard deviation of saccade end point eccentricity) for each session. Monkey G showed greater endpoint variability than Monkey C during both electrical and optogenetic stimulation (Fig. 3G and Fig. 3I, respectively). In both toward-stimulation and away-stimulation condition, electrical stimulation trials had significantly lower endpoint scatter compared to optogenetic trials in Monkey G (Fig. 3G & 3H) and in Monkey C (Fig. 3I & 3J). As expected, endpoint variability was substantially lower for away-stimulation trials than for towards-stimulation trials in both Monkey G (Fig. 3H) and Monkey C (Fig. 3J).

**Fig. 3.**
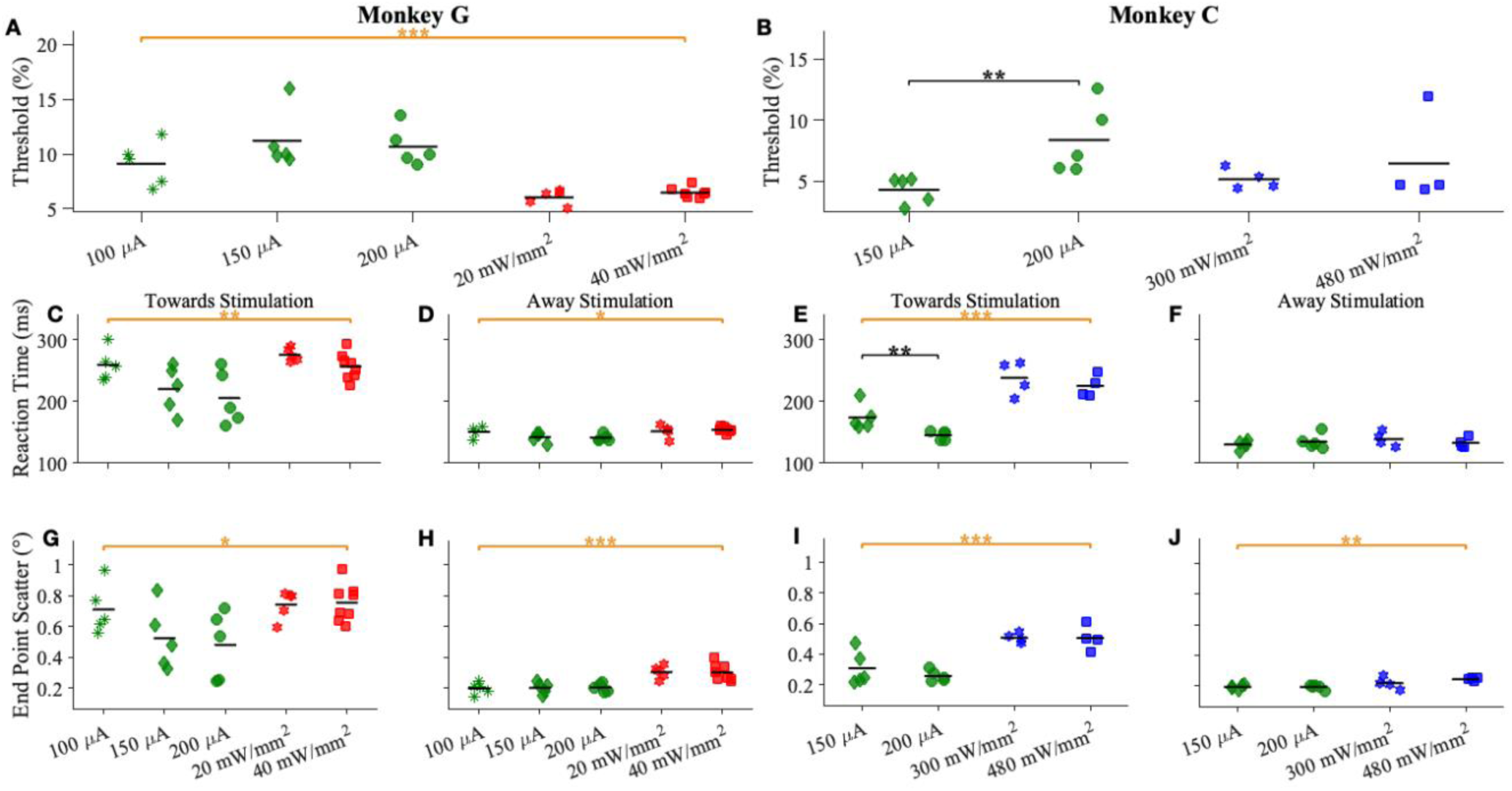
Contrast threshold, reaction time and end point scatter for electrical and optogenetic stimulation. (A and B) Contrast threshold defined as the probe contrast at which saccades were equally likely to be towards the RF and outside the RF for each current amplitude and irradiance, for Monkey G (A) and Monkey C (B). (C and D) Reaction times for saccades towards the stimulation side (C) and towards the probe side (D) across current amplitudes and irradiances, for Monkey G. (E and F) Same as (C and D) for Monkey C. (G and H) Standard deviation of saccade endpoints towards the stimulation side (G) and towards the probe side (H), for Monkey G. (I and J) Same as (E and F) for Monkey C. Stimulation conditions (current amplitudes and irradiances) are as in Fig. 2. Statistical comparisons were performed at two levels: black horizontal lines indicate significant pairwise differences between current or irradiance levels within the stimulation techniques (two-sided Wilcoxon rank-sum test, Holm-Bonferroni corrected within each panel); orange horizontal lines indicate significant differences between pooled electrical and pooled optogenetic stimulation trials (two-sided Wilcoxon rank-sum test, evaluated independently). *p<0.05, **p<0.01, ***p<0.001.

### Electrical and optogenetic stimulation induce phosphenes of comparable sizes, consistent with stimulated eccentricities

Following characterization of phosphene contrast, we characterized the perceived size of phosphenes induced by electrical and optogenetic stimulation. Results from statistical assessment are summarized in Table 1. As in the contrast discrimination task (Fig. 2A), animals performed a visual size discrimination task (Fig. 4A). Fig. 4B and 4C show the proportions of saccades directed to the RF location as a function of percentage difference between probe and standard stimulus diameter in Monkey G and C, respectively. Fig. 4E and 4F show the proportion of saccades directed to electrical or optogenetic stimulation delivered at the RF location, as a function of probe size in degrees of visual angle (dva, °) in Monkey G and C respectively.

**Fig. 4.**
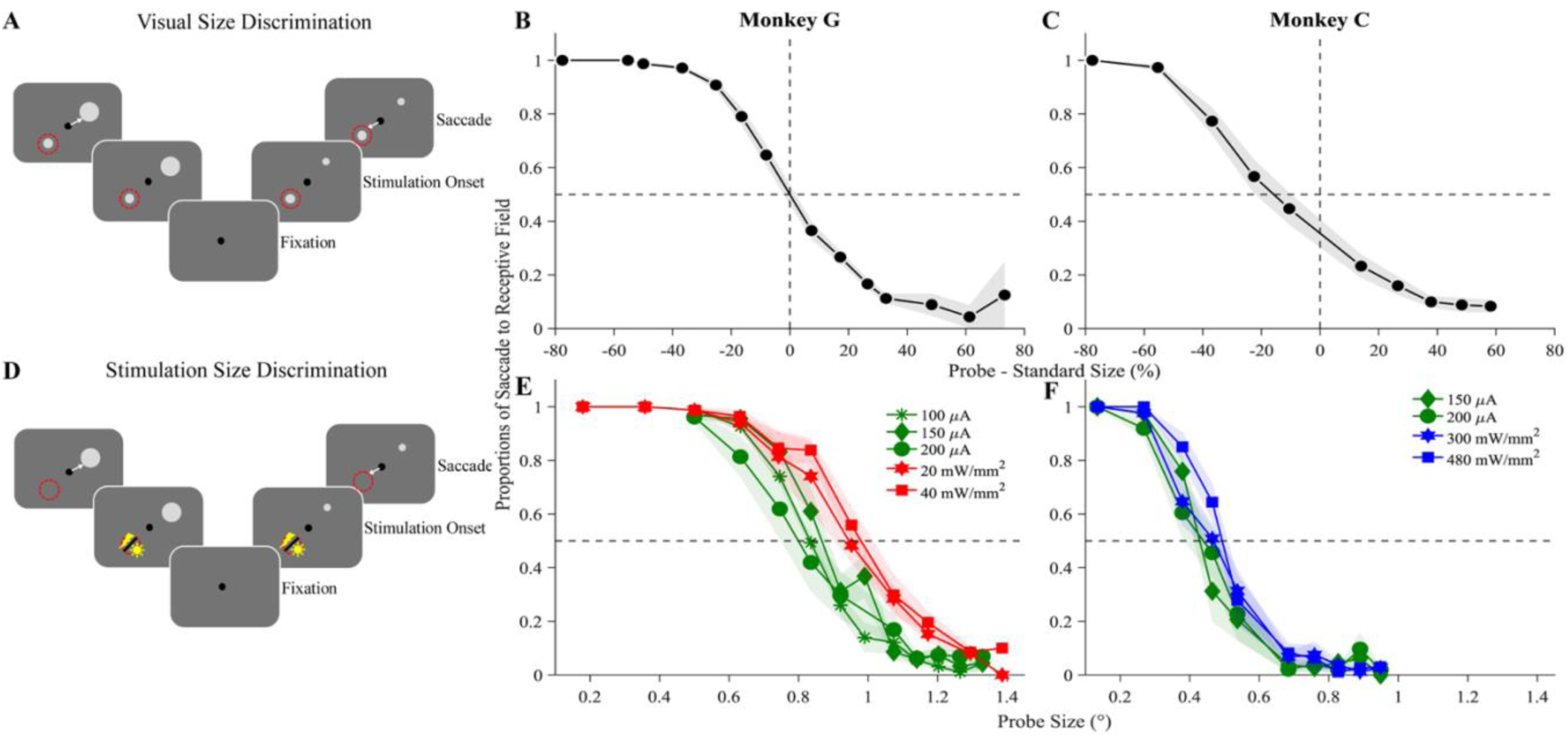
Size discrimination with visual, electrical and optogenetic stimulation. (A) Schematic of the size contrast discrimination task. On each trial, the animal saccades to the larger stimulus either in the RF or outside the RF. (B and C) Proportion of saccades towards the RF location as a function of percentage diameter difference between probe and standard stimuli for Monkey G (B) and Monkey C (C). (D) Schematic of the stimulation size discrimination task. As in Fig. 2D, the standard visual stimulus was replaced by electrical or optogenetic stimulation delivered at the RF location. (E and F) Proportion of saccades towards the RF location as a function of probe size for Monkey G (E) and Monkey C (F). Stimulation parameters are as in Fig. 2E and F.

The phosphene size threshold was defined as the probe diameter at which the monkeys were equally likely to saccade towards the RF (towards stimulation) or towards probe. Phosphene size elicited by optogenetic stimulation was larger than those elicited by electrical stimulation in Monkey G (Fig. 5A). Optogenetic stimulation elicited phosphenes of comparable perceived size to those produced by electrical stimulation in Monkey C (Fig. 5B). Perceived phosphene size did not vary significantly with current amplitudes or irradiance within either stimulation technique. Reaction times were significantly shorter for electrical than optogenetic stimulation overall in Monkey G (towards stimulation, Fig. 5C, and away stimulation, Fig. 5D). Similarly, end point scatter differed significantly between electrical and optogenetic stimulation overall in Monkey G towards stimulation (Fig. 5G) and away stimulation (Fig. 5H). We also observed a positive correlation between phosphene size and RF eccentricity (Fig. 6). The larger phosphene size observed in Monkey G may reflect optogenetic stimulation being performed at higher eccentricities than electrical stimulation.

**Fig. 5.**
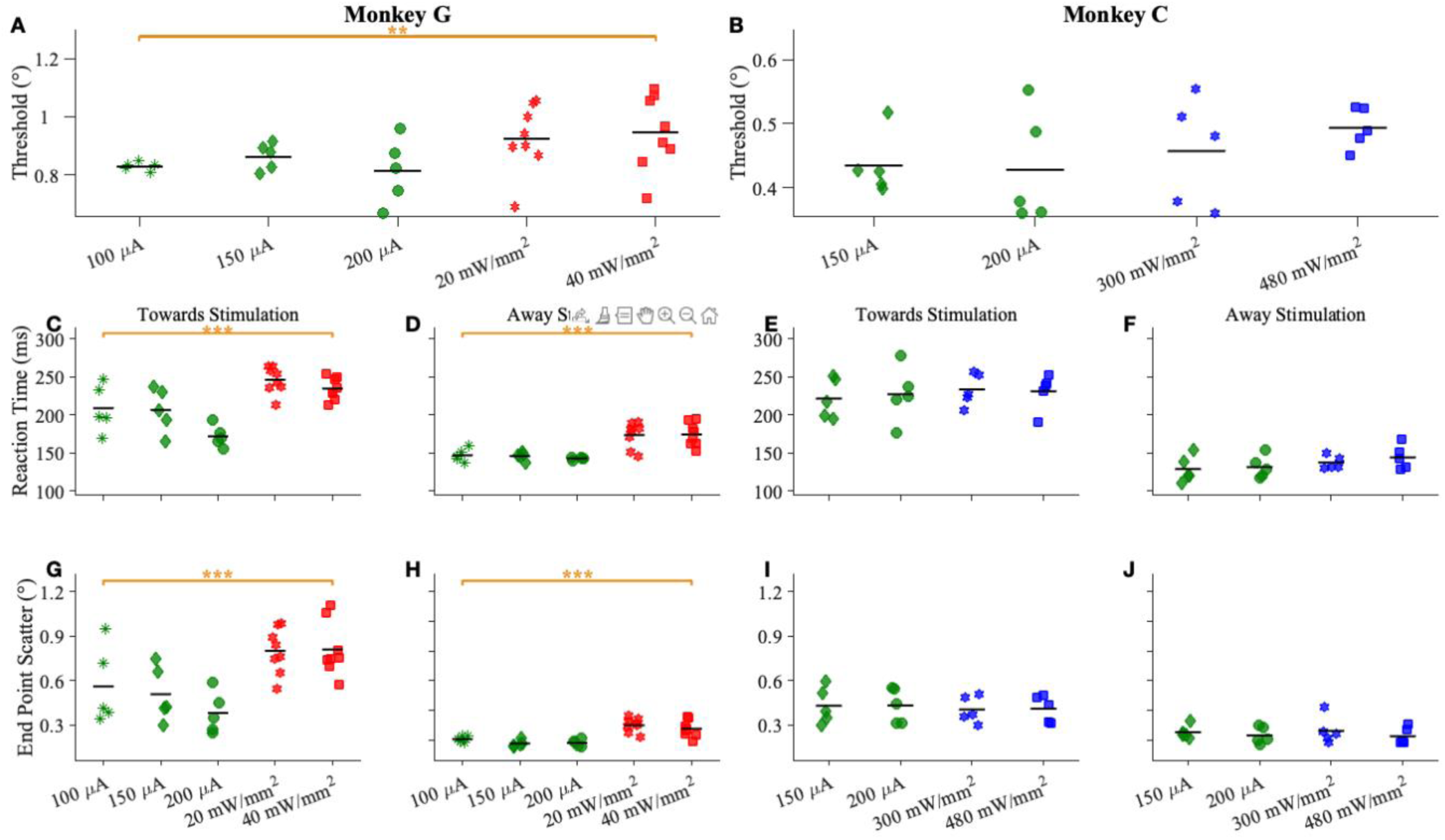
Size reaction time and end point scatter for electrical and optogenetic stimulation. Panels, stimulation conditions and statistical conventions are as in Fig. 3, but for the size discrimination task (Fig. 4). (A and B) Size threshold defined as the probe diameter at which saccades were equally likely to be towards the RF and outside the RF for Monkey G (A) and Monkey C (B). (C-F) Reaction times, (G-J) saccade endpoint standard deviation, organized as in Fig. 3.

**Fig. 6.**
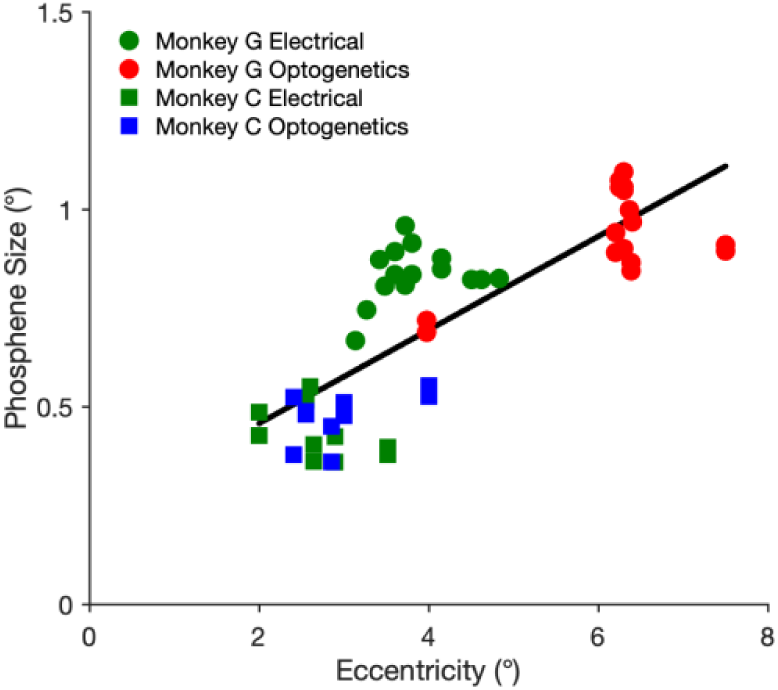
Phosphene size threshold as a function of RF eccentricity. Phosphene size threshold pooled across all sessions from the size discrimination task (Fig. 4A and B) plotted as a function of eccentricity of the recorded RF. Green points indicate electrical stimulation sessions; red and blue indicate optogenetic stimulation sessions for Monkey G and Monkey C respectively. Circles denote Monkey G and squares denote Monkey C.

## Discussion

We compared the detection probability of phosphenes elicited by electrical and optogenetic stimulation and further characterized the perceptual features of these phosphenes in the same animals. We benchmarked these phosphene related measures against perceptual readout from visual stimulation. Our results showed that electrical microstimulation reliably drove phosphene perception (d’>0) across nearly all current amplitudes tested whereas optogenetic stimulation was successful in driving phosphene perception in only a subset of sessions (48-63%). Both stimulation techniques induced phosphenes whose perceived location closely matched the RF of the stimulated site. Electrical stimulation elicited phosphenes of either higher or comparable perceived contrast than optogenetic stimulation. Phosphene size elicited by optogenetic stimulation was either larger or comparable to those induced by electrical stimulation, indicating that optogenetic stimulation does not confer a spatial resolution advantage.

Our characterization of phosphene contrast and size extends on earlier work (30) that quantitatively reported phosphene parameters induced by electrical microstimulation in non-human primate. At RF eccentricities comparable to those in Monkey G, they reported smaller phosphene sizes, potentially reflecting the lower stimulation current amplitudes used. Even though they stimulated in scotopic conditions, there Michelson contrast values are comparable to our findings. We also found a positive correlation between phosphene size and eccentricity, consistent with earlier observations in epileptic human subdural electrodes recordings (9) and the delay field measurements in macaque V1 (31). These human recordings also revealed that phosphene size increases with stimulation current but saturates at higher current levels (9), a trend also evident in our electrical stimulation results but not observed in optogenetic stimulation. We also observed a higher reaction time and end point scatter for optogenetic compared to electrical stimulation which may suggest a longer processing time and wider spread of phosphene respectively. We note that the spatial activation extent and density of viral transfection, which we did not measure, could independently influence the apparent perceived size and contrast. This also limits our direct comparison of electrical and optogenetic stimulation.

Previous work in NHP demonstrated robust optogenetic activation of V1 neurons can be associated with a wide set of behavioral effects, ranging from no reported effect, over modulation of a visual percept, to robust responses to the stimulation itself (1–3, 32–34). In these latter studies, V1 activation produced a localized visual sensation that engaged the oculomotor system and drove saccades to the stimulated locations. However, a significant confound in this paradigm is that a saccade directed at the stimulation site could reflect a reflexive oculomotor response to stimulation itself, rather than a conscious percept of a phosphene. In addition, even if a phosphene was perceived, its perceptual qualities, like size or contrast, remain unknown. Our task addresses this gap by requiring animals to respond to a target that was spatially distinct from the RF and to explicitly report trials in which no phosphene was perceived. While recent research (2) achieved high detectability with saccade probability nearing 100% at 1mW/mm^2^ irradiance level, we found that phosphene detection was lower despite using much higher irradiances. This discrepancy may reflect the use of mesoscale microLED stimulation and the highly light-sensitive opsin, ChRger in that study, but may also be related to task design. V1 optogenetic stimulation has also been shown to act as a perceptual mask, elevating detection thresholds for co-presented visual targets (34). However, as their task required animals to ignore the optogenetic stimulation and to report visual targets instead, it did not yield a direct estimate for perceiving optogenetic stimulation itself. Finally, optogenetic V1 stimulation has been demonstrated to faithfully engage V1’s computational circuits, yet without assessment of behavioral benefits. The literature on V1 thus appears in line with a considerable body of evidence indicates that optogenetic stimulation can evoke robust neuronal spiking in primate cortex while producing only weak or unreliable changes in behavior (35).

Our study also highlights the individual-animal and opsin-dependent variability that remains an issue with non-human primate optogenetics. This variability was apparent in behavioral performance, with monkey G, expressing ChrimsonR and stimulated via a 1.5 mm-diameter epidural LED detected phosphenes more reliably than monkey C, expressing ChR2 and stimulated via a 50 mm laser fiber placed inside the cortical tissue. These stimulation conditions differed across two physical measures: peak irradiance was approximately one order of magnitude higher for the laser, whereas the LED likely illuminated a much larger cortical area (Table 2). Consequently, Monkey G received lower peak irradiance but a broader activation, which was extended in depth by the deeper tissue penetration of the longer-wavelength light used to excite ChrimsonR. This pattern suggests that the spatial extent and volume of activated cortex, not only peak stimulation intensity, is an important determinant of behavioral detectability. A larger, more salient phosphene is easier to report, leading to lower contrast thresholds and to overall better detectability. This prediction is consistent with our observation in monkey G. However, because the opsin, individual animal, light source and illumination geometry all co-vary in our two-animal study, these confounding factors cannot be isolated. Disentangling these variables would require a within animal comparison matching both the illuminated area and irradiance across different opsins.

**Table 2:**
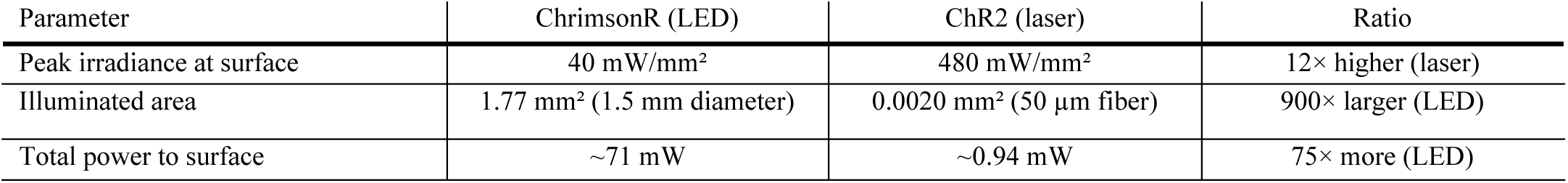
Light delivery differed between the two opsin/light-source configurations.

These open questions carry significant implications for the design of optogenetic based neuroprosthesis, for which detection reliability not merely the presence of a behavioral effect, would serve as a primary benchmark. While electrical microstimulation can produce phosphenes reliably, its comparatively limited long-term biocompatibility (34) makes optogenetics still a compelling alternative. The neural mechanisms underlying the differences in perceptual detectability we observed, however remained unresolved. Directly comparing the pattern of neural activation evoked by electrical versus optogenetic stimulation will be an important next step towards understanding the basis of these differences in detectability and discriminability.

## Supplementary Figures

**Fig. S1:**
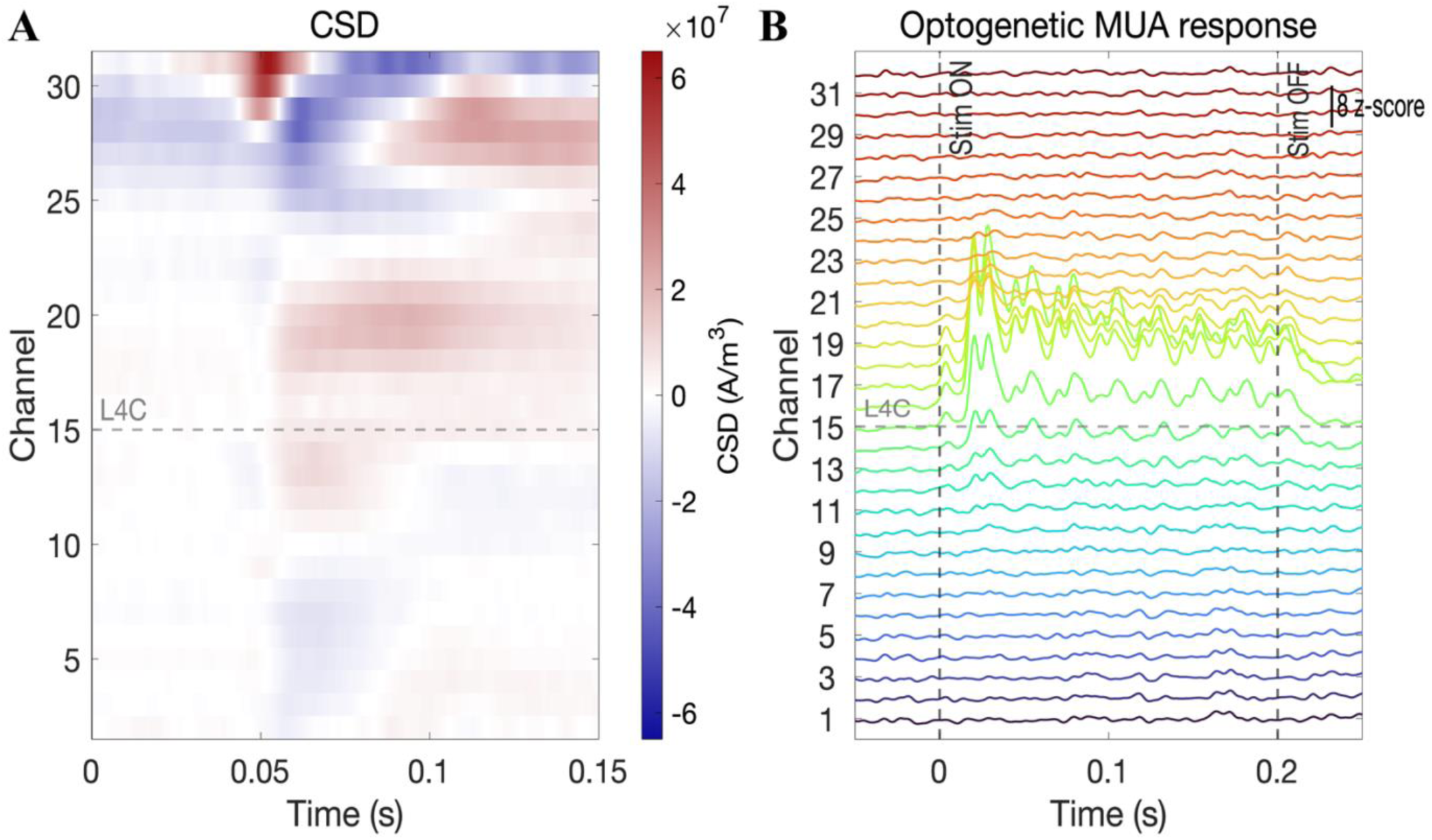
Electrophysiological confirmation in Monkey. **C** (A) CSD profile for an example recording site in Monkey C during an optogenetic session. (B) MUA activity from the same session, evoked by invasive blue laser optogenetic stimulation.

**Fig. S2:**
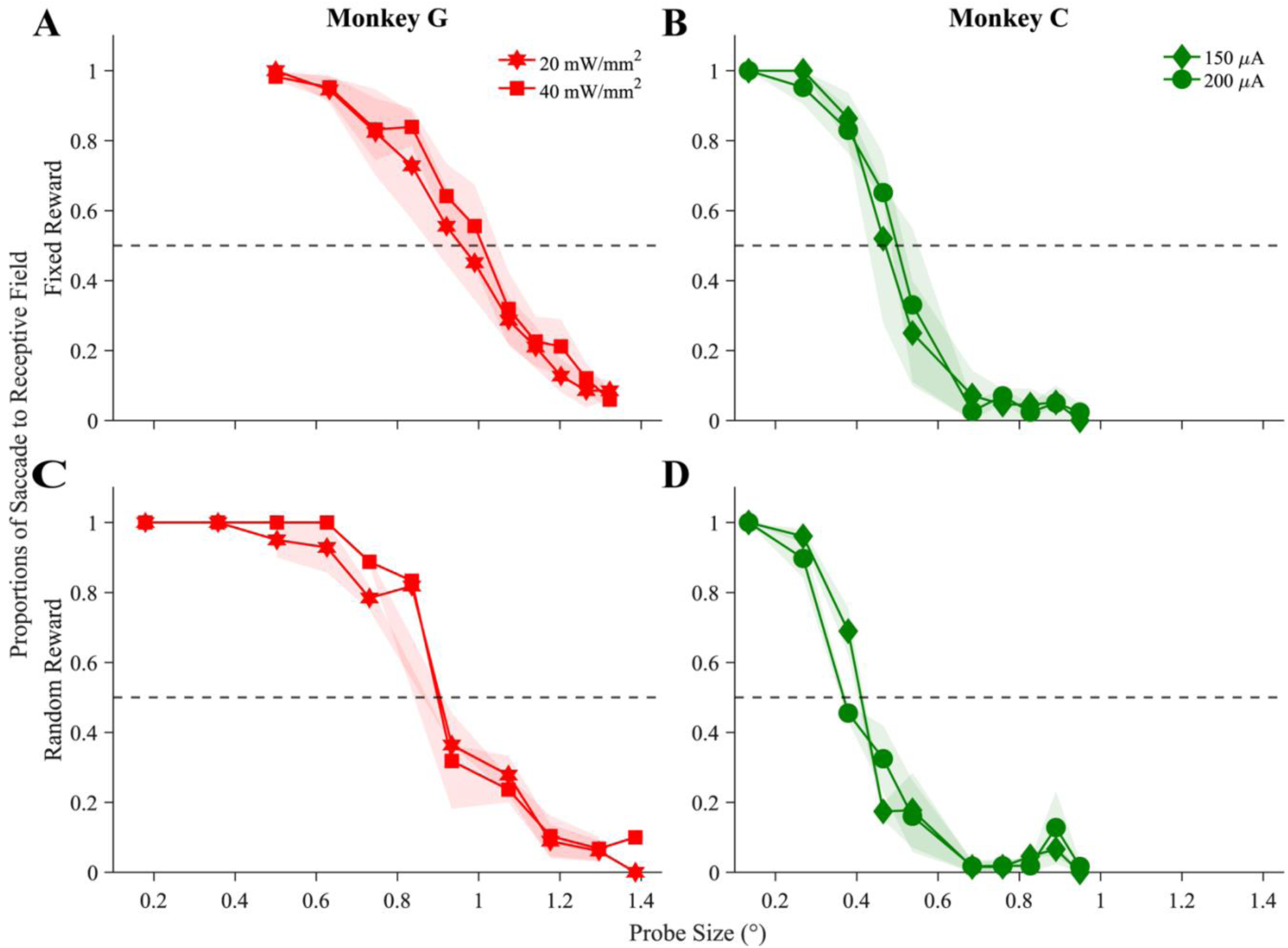
Size discrimination with optogenetic and electrical stimulation under fixed and random reward methods. Proportion of saccades towards the RF location as a function of probe size (A) Monkey G, optogenetic stimulation sessions, fixed reward. (B) Monkey C, electrical stimulation sessions, fixed reward. (C) Monkey G, optogenetic stimulation sessions, random reward. (D) Monkey C, electrical stimulation sessions, random reward.

## Acknowledgements

We thank the veterinary and animal care staff at the University of Fribourg and the Experimental Animal Center of the University of Bern for their support with animal husbandry, anesthesia and perioperative care. We are grateful to the members of the Primate Neuroscience Platform and the Visual Cognition Lab of Prof Gregor Rainer for helpful discussions on experimental design and analysis, and to the technical and workshop staff of the Department of Neuro- and Movement Sciences for support. This work was supported by Swiss National Science Foundation, grant no. 310030_204544 and 31NE30_203974 to M.C.S. and Carigest Foundation, grant ID 22388 to D.G.

## Author contributions

M.C.S. conceived and supervised the study. K.L., M.F.-K., J.C.-V. and M.C.S. designed the experiments. M.F.-K. performed the viral injections and optogenetic experiments and analysis, K. L. performed the electrical stimulation experiments and analysis. A.B. providing veterinary and anesthesia support. J.G., F.L. and M.H. performed animal habituation and training. J.C.-V. developed the stimulation and recording hardware and software. D.G. contributed to the design of the electrical stimulation approach and co-supervised the study. K.L., M.F.-K. and M.C.S. wrote the manuscript, with input from all authors.

